# predPPI-GReMLIN: prediction of protein-protein interactions through mining of conserved bipartite graphs

**DOI:** 10.64898/2025.12.03.692055

**Authors:** Moruf A. Adeagbo, Valdete M. Gonçalves-Almeida, Sandro C. Izidoro, Sabrina A. Silveira

**Affiliations:** Department of Computer Science, Universidade Federal de Viçosa, 36570-900, Minas Gerais, Brazil; Department of Computer Sciences, Abiola Ajimobi Technical University, Ibadan, 200255, Nigeria; Computer Center, Campus Almenara, Instituto Federal do Norte de Minas Gerais, 39900-000, Minas Gerais, Brazil; Institute of Technological Sciences, Universidade Federal de Itajubá, 35903-087, Minas Gerais, Brazil

**Keywords:** protein-protein interactions, bipartite graphs, graph mining, machine learning, structural bioinformatics

## Abstract

Protein–protein interactions (PPIs) play a central role in elucidating cellular mechanisms. However, a substantial gap remains in current prediction models, as they frequently overlook the structural and physicochemical context governing molecular binding, thereby limiting predictive accuracy. To address this limitation, we introduce predPPI-GReMLIN, a graph-based framework that represents protein–protein interfaces as bipartite graphs, integrating atomic-level physicochemical descriptors, spatial distance constraints, and conserved substructure mining to perform PPI prediction. The method employs a graph-search strategy to detect interface interaction patterns and conducts ligand swapping with complexes that share the same patterns as the query, enabling the prediction of novel interaction partners. Evaluations across multiple datasets—including CAMP (protein–peptide), Yeast (binary PPI), and TAGPPI (multi-class PPI)—demonstrate consistently strong predictive performance, achieving precision, recall, accuracy, and F1-scores exceeding 97% on binary classification benchmarks and surpassing state-of-the-art sequence- and structure-based approaches. Furthermore, incorporating solvent-accessible surface area (SASA)-derived features improved multi-class interaction-type classification accuracy to 57.74%. A case study on the SARS-CoV-2 spike–ACE2 complex further validated the approach: docking simulations using ligands predicted by our method reproduced native-like binding energetics comparable to redocking results. Additionally, a comparison between docking using a predPPI-GReMLIN-predicted ligand and docking using ligands selected at random from the PDB yielded a highly significant p-value (*p* = 5.1 × 10^−9^), indicating a robust statistical difference in binding performance. Collectively, these findings demonstrate predPPI-GReMLIN’s ability to capture conserved structural determinants of PPIs, providing a robust and interpretable framework for protein interaction prediction and ligand discovery. The dataset and source code for the experiments are publicly available at https://github.com/morufwork/predPPIGReMLIN.git.

## Introduction

Proteins are macromolecules that are an essential part of all organisms, composed of one or more chains of amino acids playing crucial roles in various biological processes [37, 9]. Their functions in the body are diverse and extend beyond repairing tissues, regulating chemical reactions, supporting growth, and aiding the transportation of molecules [30, 44]. Proteins interact with each other through specific binding sites, using complementary shapes, and surface charges, among other factors, to drive these interactions [31, 50]. Protein-protein interactions can form complexes that carry out various cellular processes, from catalyzing biochemical reactions to regulating signal transduction pathways and maintaining intracellular architecture [54].

The Protein Data Bank (PDB) [17, 13, 39] is an extensive repository that contains the 3D structures of proteins that are freely available to researchers. The PDB includes over 200,000 experimentally determined 3D protein structures and nucleic acids solved using methods such as X-ray crystallography, Nuclear Magnetic Resonance (NMR) spectroscopy, and and cryo-electron microscopy. With ongoing advances in technology and structural biology, the PDB is regularly updated with new protein structures. Although these experimental approaches have provided numerous protein structures, they are often time-consuming and resource-intensive [41].

AlphaFold [29, 45, 20, 1, 46], an AI-powered tool, has revolutionized structural biology by accurately predicting protein tructures from amino acid sequences. In addition to the experimental data found in the PDB, the AlphaFold Protein Structure Database now contains more than 214 million predicted protein structures [46]. Researchers can now anticipate the structures of proteins from nearly the whole human proteome and proteins from various other species, even those previously deemed unsolvable by experimental methods. AlphaFold makes it easy to quickly and efficiently predict the structures of proteins [36]. Researchers can use these predictions to improve the design of molecules that interact with proteins, which enhances drug effectiveness and shortens development time.

Several computational approaches have been proposed to predict PPIs, leveraging recent advances in deep learning, graph-based models, and structural biology. Notably, the availability of high-quality structural predictions from AlphaFold has opened new avenues for incorporating 3D conformation into PPI modeling.

Sequence-driven models dominate many recent studies due to their ease of use and availability of large-scale data. The ProtBert-BiGRU-Attention model was employed for comparison and ablation analyses, demonstrating strong predictive performance and accuracy in PPI prediction [33]. The ProtBert-BiGRU-Attention model demonstrates robust predictive accuracy; nonetheless, it predominantly depends on protein sequence data, which may overlook essential structural and physicochemical attributes that affect interactions. Similarly, CAMP introduces a sequence-based deep learning framework for predicting peptide-protein interactions (PPIs) and identifying binding residues [23]. However, the extraction of features heavily relies on sequence-based profiles, ignoring explicit atomic interactions that drive peptide-protein binding, while CNN-based architectures, though effective for local feature extraction, inherently struggle to capture long-range dependencies in protein structures.

To overcome the limitations of purely sequence-based models, hybrid strategies have emerged. KSGPPI predicts better PPI by using ESM-2-CKSAAP and Node2vec to extract features from protein sequences and interaction networks, respectively [16]. While the model extracts features from both sequences and interaction networks, it does not explicitly consider 3D structural data that are crucial for understanding PPIs. Similarly, MultiPPIs achieved 86.03% accuracy, 93.03% AUC, and 82.69% sensitivity on PPIs by integrating features derived from protein sequences with multi-source associations network built from known relationships among proteins, drugs, miRNAs, lncRNAs, and diseases [55]. This method helps connect multiple sources, but it lacks 3D structural data, atomic-level physicochemical details, and direct modeling of binding interactions. Furthermore, [15] employed deep neural network and feature fusion to precisely predict protein-protein interactions (PPIs) in yeast and human datasets. The study successfully predicted interactions within PPI networks. This model primarily relies on sequence-based embeddings and handcrafted features. Protein interactions are inherently 3D; ignoring spatial conformations and binding site structures can lead to false positives [18, 19, 7].

Recent efforts have sought to embed structural context into PPI prediction. TAGPPI combines sequence-based convolutional learning with graph-based contact maps obtained from AlphaFold [38]. This approach improves over purely sequence-based models by incorporating spatial topology but do not incorporate physicochemical properties (e.g., charge, hydrophobicity, and electrostatics). In contrast, studies such as [25] utilize template-based modeling, docking simulations, and surface patch analysis, offering more detailed structural interpretations. However, these methods face challenges with transient or weakly bound interactions and can be computationally intensive.

FDPPI combines a quartile-based algorithm and batch normalization, achieving high predictive metrics (98.34% precision, 98.09% accuracy, and 97.72% sensitivity) of PPI prediction using human data [22]. However, its reliance on sequence-based data, which lacks structural and contextual depth, limits its interpretability, generalizability, and potential for providing deeper biological insight.

Another promising direction involves building a protein network and utilizing a graph neural network to extract fused features, resulting in a higher level of accuracy compared to existing models [51]. However, this study evaluated performance solely based on accuracy, which is only one of several important metrics for assessing a model’s overall effectiveness.

To address the limitations of previous studies, we introduce predPPI-GReMLIN, a graph-based strategy for predicting protein-protein interactions that incorporates atomic-level physicochemical properties, spatial distance criteria, and structural pattern matching. For each protein complex, the protein-protein interfaces were calculated at the atomic level using physicochemical properties of atoms and the distance criteria between them [12]. PPIs interface are modeled as bipartite graphs, where nodes and edges represent non covalent interactions between them. The interaction types depend on the distance criteria and the physicochemical types of nodes. This bipartite representation considers only the contacts between atoms of different protein chains. The resulting set of graphs, their conserved patterns serve as an input for protein-protein interaction prediction. Structural pattern searching techniques enables the identification of conserved structural arrangements within protein-protein interfaces, potentially aiding in ligand prediction within protein complexes. This method captures structural and spatial information, provides a more accurate representation of binding interfaces, and improves the handling of novel interactions.

## Methods

This section explains predPPIGReMLIN, our graph based strategy to predict protein-protein interactions. We explain the problem modeling, the datasets used in the experimental evaluation and the evaluation strategy.

### predPPI-GReMLIN

The starting point for this work is a graph based strategy named ppiGReMLIN, proposed by our group in [34] to identify conserved structural arrangements in protein-protein interfaces. The original ppiGReMLIN comprises three main components: *Data Acquisition and Modelling, Clustering Analysis*, and *Conserved Substructure Mining*. In the *Data Acquisition and Modelling* module, interactions at the protein–protein interfaces are computed at the atomic level for a set of complexes, based on the physicochemical properties of atoms and distance criteria. These interactions are modeled as bipartite graphs as illustrated in the Figure 1. The *Clustering Analysis* module receives as input a collection of graphs representing protein–protein interfaces and partitions them into clusters of structurally similar graphs, in preparation for frequent substructure identification in the subsequent module. These clusters are then passed to the *Conserved Substructure Mining* module, where a frequent subgraph mining (FSM) task is performed to identify conserved substructures within each group.

**Fig. 1.**
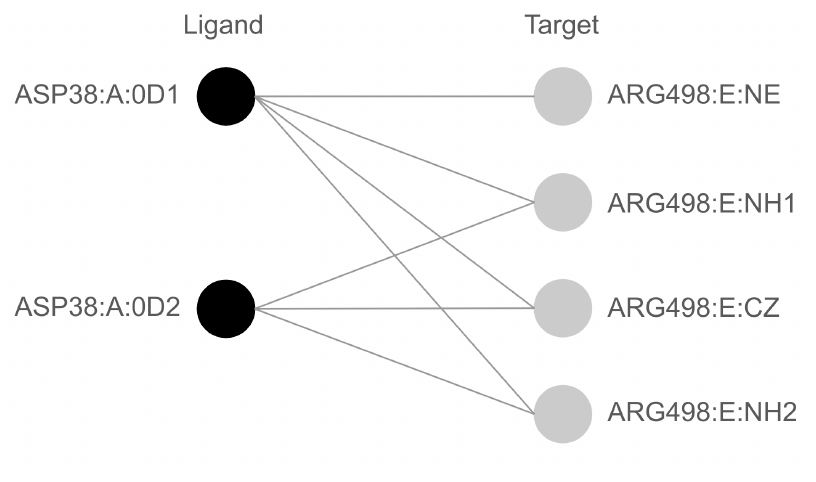
A bipartite graph representing the complex interface of PDB ID 7BH9 (SARS-CoV-2 RBD-62 in complex with the ACE2 peptidase domain). The atom-level bipartite interface graph illustrates the interactions between the ligand (ACE2, ASP38 in chain A) and the target (SARS-CoV-2 RBD, ARG498 in chain E). Black nodes represent atoms on the ligand side, whereas grey nodes represent those on the target side. Edges denote non-covalent interactions between them.

The results demonstrate that ppiGReMLIN is effective in automatically identifying conserved structures previously described in the literature. Precision ranged from 69% to 100%, with a recall of 100% when considering relevant structures reported in the literature for the datasets considered.

In the present work, we extend the previously proposed computational strategy to perform protein-protein interaction prediction, which is an important step towards ligand prediction for targets of interest (predictive ppiGReMLIN or predPPI-GReMLIN). Both the ligands and the targets are protein-based. Figure 2 presents the predPPIGReMLIN workflow and was inspired in [32]. It is segmented into three mains steps and an optional step which is represented by ppiGReMLIN. There are 2 options for the *Input* step: (i) a set of complexes (bipartite graphs) resulting from ppiGReMLIN conserved substructure mining from a set of proteins; or (ii) a set of user selected protein complexes which will have their interface modeled as bipartite graphs. Next we have the *Pattern search* step and finally the *Ligand prediction* step.

**Fig. 2.**
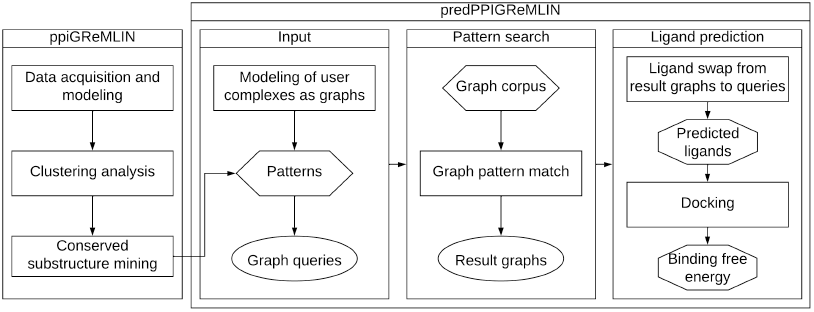
predPPIGReMLIN workflow. The workflow is segmented in three main steps, with ppiGReMLIN being an optional step to provide input data: *Input, Pattern search* and *Ligand prediction*. Rectangular boxes represent action steps, hexagonal boxes depict input data or parameters, ellipsoids are intermediate data and octagonal boxes represent output data.

Prediction is carried out by predPPI-GReMLIN as follows: the conserved structural arrangements in protein–protein interfaces—i.e., the patterns identified—are used to predict ligands for proteins of interest. The bipartite graphs representing these patterns consist of one set of nodes labelled as ligand and the other set labelled as target. There are no interactions between nodes on the ligand side nor on the target side. A search for such patterns, which we named *query graphs*, is conducted across the universe of protein–protein complexes (modeled as interface graphs), which we named *Graph corpus*, to identify those that contain the same patterns (graphs) at their interfaces. When a match to a pattern is found, the underlying hypothesis is that the molecule associated with the ligand side of the matched bipartite graph (pattern) constitutes a potential ligand for the molecule on the target side of the query graph. Figure 3 illustrates how the graph pattern search is performed and how its result provides potential ligands that can be swapped with the ligand from the query, enabling the prediction of potential ligand-binding partners for the target interface.

**Fig. 3.**
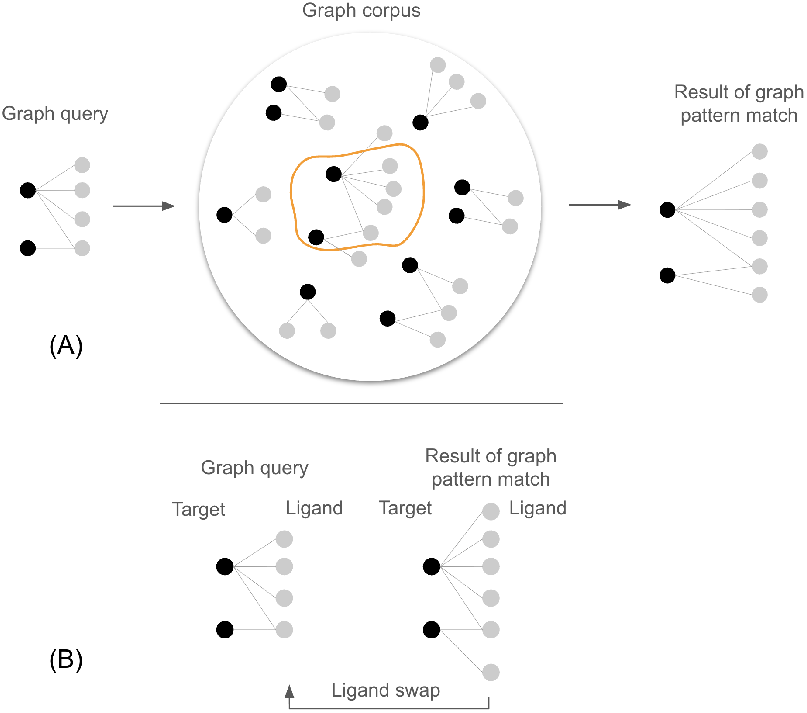
Graph pattern search. In (A), a *query graph* (representing an interface pattern of interest) is searched against a universe of protein–protein complexes, referred to as the *graph corpus*, to identify those graphs that contain the same pattern as the *query graph*. Our hypothesis is that the molecule associated with the ligand side of the graphs in the *graph pattern match result* is a potential ligand for the molecule on the target side of the *query graph*. In (B), the process by which a ligand swap can potentially be performed—from the original ligand in the *query graph* to the predicted ligand obtained from the *graph pattern match result* —is illustrated.

#### Evaluation strategy

To compare predPPGReMLIN with other methods the process begins by constructing interface graph representations of protein-protein interactions from a *training dataset*. Each protein complex is modeled as a bipartite graph based on physicochemical properties and distance criteria. These graphs serve as *graph queries*. Similarly, we compute *interface graphs* for the collection of all known protein complexes experimentally solved, which are available from PDB [2, 5]. These *interface graphs* from PDB were named as *graph corpus*.

The *graph queries* are matched against the *graph corpus* to find the same graphs, representing potential interaction patterns. This graph matching step produces a set of graphs which we named *graph pattern match*.

Finally, an *interface graph* is computed for each protein complex of the *test dataset*. These graphs serve as *graph tests*. We match the *graph tests* against the result of *graph pattern match* and evaluate the predPPI-GReMLIN results using standard classification metrics such as precision, recall, accuracy, F1 score and Matthews Correlation Coefficient (MCC).

### Data

This study used entries from the RCSB Protein Data Bank (PDB) [2, 5], that involves protein-protein or protein-peptide complexes to compose the the *graph corpus*. The dataset includes experimentally determined protein complexes from various structural determination methods, such as X-ray crystallography, cryo-electron microscopy, and NMR spectroscopy.

To benchmark and evaluate predPPGReMLIN’s quality, generality and applicability in real-world, we incorporated four datasets with specific purposes:

1. CAMP dataset: composed by 7,417 positive peptide-protein interaction pairs involving 3,412 unique proteins and 5,399 unique peptides. Among these, 6,581 pairs have residue-level binding labels. Negative pairs were generated by randomly shuffling non-interacting protein-peptide pairs. For our study, we utilized the training and test datasets provided in Supplementary Tables 12 and 13 of CAMP [23]. The purpose of this dataset is to compare our method with CAMP. This dataset is employed to assess the ability of our proposed strategy to predict protein–protein interactions from complex structures. The performance on peptide–protein interactions will further serve to validate predPPGReMLIN’s effectiveness in capturing structural interaction patterns for broader applicability.
2. SARS-CoV-2 dataset: includes experimentally solved complexes of receptor-binding domain (RBD) of the spike protein of SARS-CoV-2 bound to the cell receptor ACE2. The dataset covers multiple variants, including Orig, Alpha, Beta, Delta, Gamma, Kappa, Omicron, and structures without a specific variant annotation (None), with resolutions < 3.0 Å (43 structures). This dataset is used to raise candidate ligands for the RBD of the spike protein of SARS-CoV-2, using the predPPIGReMLIN, and validate them via a docking protocol.
3. Yeast sequence dataset: includes 2,497 protein sequences with 5,594 positive and 5,594 negative interaction samples of the yeast dataset used in the method PIPR[6]. This dataset is part of Guo’s benchmark collection [4] in which positive samples are known interacting protein sequence pairs. Yeast dataset has become a standard benchmark, commonly used in the evaluation of state-of-the-art methods for PPI prediction, as the positive interactions are experimentally validated. This dataset is used to compare predPPIGReMLIN with a variety of methods.
4. Multi-class dataset: comprises 16,278 proteins and 75,875 protein-protein interaction samples from TAGPPI [4], annotated with one of seven interaction types: activation, binding, catalysis, expression, inhibition, post-translational modification, and reaction. The dataset is composed by Homo sapiens protein structures generated by AlphaFold, and its interaction labels were obtained by aligning these proteins with interaction data from the STRING database [43]. The purpose of this dataset is to compare our method with TAGPPI in the task of predicting different protein interaction types.

## Results and discussion

To assess the predictive capability of our method for predicting PPI, we conducted a comprehensive set of experiments. First, we compare predPPI-GReMLIN with other four high quality PPI prediction methods. Next, we compare our strategy with 10 sequence based methods. Then, we present comparative results regarding methods capable to predict the type of PPI. Finally, we apply our method in the prediction of ligands for the RBD of the spike protein of the Sars-CoV-2, comparing the binding free energies of the predicted ligands docked with receptors with those of real complexes available at the PDB, through a redocking protocol, and with ligands chosen at random.

### predPPI-GReMLIN compared with state-of-the-art methods

We evaluated the performance of predPPI-GReMLIN on the CAMP dataset. This evaluation aims to accurately assess the model’s ability to predict binary protein–peptide interactions using structural information. We obtained the chains of protein complexes from the *training* and *test* datasets provided by CAMP. A *corpus* dataset was created based on all PDB complexes involving protein–protein or protein–peptide interactions. Each complex from these datasets was modeled as an interface graph. The three types of graphs constructed are:

- *Graph corpus D*(*G*): 115, 225 graphs derived from protein complexes in the PDB used as the structural reference base.
- *Graph queries Q*(*G*): 8, 486 graphs representing interaction patterns from the *training dataset*.
- *Graph test T* (*G*): 1, 218 graphs constructed from the *test dataset*.

Based on the *Evaluation strategy* described in *Methods*, we performed a graph-based pattern matching between the *graph queries* and *graph corpus*, which resulted in a set named *graph pattern match*. The *graph pattern match* represents all *actual positives*, and the graphs from *graph corpus* that do not match *graph queries* represent all *actual negatives*. Then the *test graphs* are matched against all actual positives to generate true positive and false negative patterns. Similarly, the *test graphs* were matched against all *actual negatives* to generate true negative and false positive patterns. Figure S1 explains the process in detail in the supplementary material. We evaluate the result using precision, recall, accuracy, F1 score, and MCC. predPPI-GReMLIN shows 99.82% precision, 97.26% recall, 97.23% accuracy, 98.52% F1-score, and 0.79 MCC.

To support machine learning analysis for binary classification of graph patterns generated by predPPI-GReMLIN, we designed a comprehensive pipeline for extracting structural, topological, and biochemical features from the graph patterns. All *actual positive* and all *actual negative* graphs are labelled as 1 and 0, respectively. To ensure a consistent feature space across all graph patterns, we first constructed global vocabulary metadata by collecting all unique names of residue, node, and edge across the complete set of graph patterns (all actual positive and negative). These vocabularies are used along with other metadata in node and edge features. We also include structural properties, centrality measures, and topological metrics in the graph features. To ensure no data leakage, all graph features are extracted independently, without reference to class labels. Each graph was encoded as a feature vector. This pipeline produces a structured, interpretable dataset of graph features that enables the application of machine learning models to biomolecular structures. A summarized breakdown of these feature vectors is provided in Figure S2 of the Supplementary Material.

We employed a threshold-based cluster-level stratified cross-validation strategy to ensure stability and robust generalization of the Random Forest classifier across varying structural groupings of the feature space. We apply K-Means to partition the entire feature space into clusters at multiple thresholds from 0.3 to 0.6 (in steps of 0.1). Each cluster was independently processed and subjected to 5-fold Stratified Cross-Validation using StratifiedKFold to ensure balanced representation of both classes within each fold. We trained using a Random Forest classifier on each fold and evaluated on the corresponding test fold. After aggregating the predictions across all clusters and folds for a given threshold, we compute ROC-AUC and PR-AUC scores to quantify the classifier’s ability to distinguish between the two classes. We tracked the ROC-AUC and PR-AUC across all threshold experiments to identify the best-performing clustering configuration. Figure S3 in the supplementary material illustrates how the clustering threshold affects the model performance.

The result shows that ROC-AUC remains above 0.998 across all tested thresholds, but begins to decline at threshold 0.6. PR-AUC mirrors this trend with high stability up to threshold 0.5 and a slight decrease at 0.6. However, PR-AUC score consistently above 0.998 confirms robust precision even under varied clustering conditions. Overall, the results reveal that the classifier maintains consistently high ROC-AUC and PR-AUC scores across thresholds, indicating strong generalization within structurally defined subgroups of the data.

We compared our results with those of existing methods. Figure 4 shows that ROC-AUC and PR-AUC across the thresholds surpassed baseline techniques such as the similarity-based matrix factorization method (NRLMF) [27], PIPR [6], the deep learning-based model for CPI prediction (DeepDTA) [56], and CAMP [23].

**Fig. 4.**
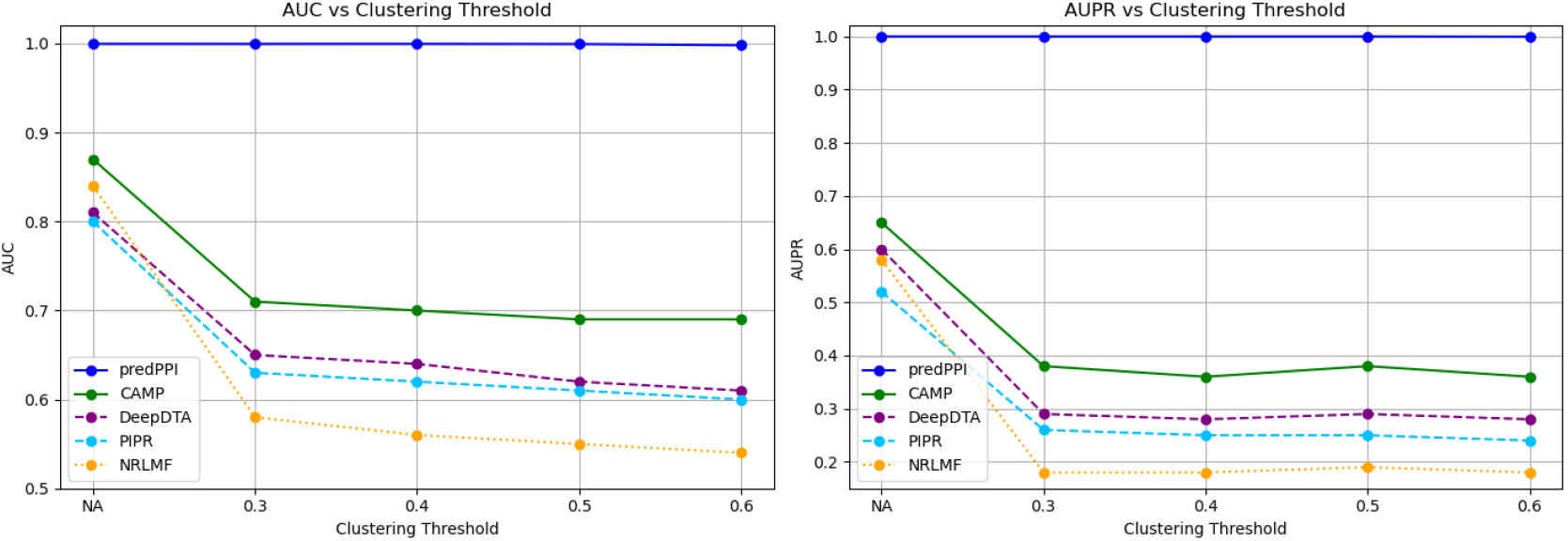
Performance comparison of prediction models across clustering thresholds. AUC (left) and AUPR (right) of five models (predPPI, CAMP, DeepDTA, PIPR, and NRLMF) evaluated at clustering thresholds 0.3, 0.4, 0.5, and 0.6. predPPI maintains high performance, while other models decline with increasing thresholds.

### predPPGReMLIN compared with sequence based methods

As previously mentioned, there is a considerable number of sequence based computational methods for PPI prediction as there is much more experimental sequence data available, compared to experimentally solved structures [40, 3, 26]. predPPI-GReMLIN is a structure based method that models protein-protein interactions as bipartite graphs and predicts ligands based on conserved graph patterns. Some representative sequence based methods methods are MCD-SVM [53], RF-LPQ [47], KNN-CTD [49], EELM-PCA [52], DeepPPI [10], SAE [42], DPPI [14], DNN-PPI [24], PIPR [6], whith emphasis on TAGPPI [4], considered state-of-the-art. Therefore, a natural question that arises is how the predictions of predPPI-GReMLIN can be compared with the predictions of these methods.

We evaluated the performance of predPPI-GReMLIN using the yeast dataset (as described in the Yeast sequence dataset subsection of the Data), which comprises 5,594 positive and 5,594 negative protein-protein interaction (PPI) samples. AlphaFold Multimer [11, 21], predicted 3D complexes for each protein pair, and we filtered out those with zero interface contacts or low predicted DockQ scores. Each complex was modeled as an interface graph (as detailed in the Methods: Evaluation Strategy section) and subjected to a 5-fold cross-validation strategy. Each complex was modeled as an interface graph (as detailed in the *Methods: predPPI-GReMLIN* section) and subjected to a 5-fold cross-validation strategy.

The evaluation strategy used in this experiment follows the procedure described in the *Evaluation Strategy subsection of the Methods*. However, here the *Yeast sequence dataset* was segmented into five folds, to perform cross-validation. PDB entries involving protein-protein or protein-peptide complexes were modeled as interface graphs to compose the *graph corpus*. After segmenting *Yeast sequence dataset* into five folds and calculate their corresponding interface graphs, one fold is used to compose the *graph queries* and the remaining four folds are the *graph test*. The *graph queries* are matched against the *graph corpus* to create the the *graph pattern match*, which represents all actual positives. The graphs from *graph corpus* that do not match *graph queries* represent all actual negatives. We used the remaining four folds *graph tests* against all actual positives to generate true positives and false negatives. Similarly, the *graph tests* were matched against all actual negatives to generate true negatives and false positives. Finally, the process is repeated for every one of the five folds. Table S1 from Supplementary Material shows the overall evaluation of the predPPI-GReMLIN across five folds of cross-validation.

predPPI-GReMLIN was compared with previous sequence-based methods on the benchmark yeast dataset. Table 1 shows the performance comparison between our method and ten high quality sequence-based methods on the mentioned dataset. predPPI-GReMLIN shows the best performance across most metrics, especially accuracy, precision, sensitivity, and F1 score. The TAGPPI method is the second, particularly in MCC and Specificity, followed by PIPR. The comparison demonstrates that predPPI-GReMLIN is the leading method for PPI prediction on the Yeast dataset, outperforming other state-of-the-art methods in almost all evaluated metrics. These results demonstrate the robustness and predictive capacity of the predPPI-GReMLIN for identifying protein–protein interactions, even in the absence of experimentally resolved sequences, a limitation that may be overcome through the use of modelled structures generated by tools such as AlphaFold.

**Table 1.** The Performance Comparison between predPPI-GReMLIN and Ten State-of-the-art Sequence-based Methods.

| Method | Acc. | Pre. | Sens. | Spec. | F1 | MCC |
| --- | --- | --- | --- | --- | --- | --- |
| MCD-SVM | 91.36 | 91.94 | 90.67 | NA | 91.30 | 84.21 |
| RF-LPQ | 93.92 | 96.45 | 91.10 | NA | 93.70 | 88.56 |
| kNN-CTD | 86.15 | 90.24 | 81.03 | NA | 85.39 | NA |
| EELM-PCA | 86.99 | 87.59 | 86.15 | NA | 86.86 | 77.36 |
| DeepPPI | 94.43 | 96.65 | 92.06 | NA | 94.30 | 88.97 |
| SAE | 67.17 | 66.90 | 68.06 | 66.30 | 67.44 | 34.39 |
| DPPI | 94.55 | 96.98 | 92.24 | NA | 94.41 | NA |
| DNN-PPI | 76.61 | 75.10 | 79.63 | 73.59 | 77.29 | 53.32 |
| PIPR | 97.09 | 97.00 | 97.17 | 97.00 | 97.09 | 94.17 |
| TAGPPI | 97.81 | 98.10 | 98.26 | **98.10** | 98.18 | **95.63** |
| **predPPI-GReMLIN** | **99.31** | **99.64** | **99.64** | 93.31 | **99.64** | 92.94 |

*Note:* NA indicates that the corresponding metric is not available from the original paper. Bold font indicates the best result in each column. All results are reported in percentage (%).

### Predicting protein interaction types

In the previous experiments, the objective was to predict whether or not an interaction existed between protein chains. In this experiment, however, the aim is to predict the specific type of PPI from the following categories: activation, binding, catalysis, expression, inhibition, post-translational modification, and reaction. We employed data from the Multi-class dataset, detailed in Data subsection. As this dataset provides sequences with UniProt identifiers [8], these were mapped to the AlphaFold database, from which the protein structures used in this study were obtained. During preprocessing, we excluded sequences with missing values or those lacking valid protein structures, resulting in 15,987 (98.21%) out of 16,278 proteins and 67,578 (89.06%) out of 75,875 samples being retained for further analysis. The mapping between UniProt and AlphaFold identifiers is presented in Supplementary Material Table S2.

Here proteins were modelled in a different manner from the previous experiments. Each complete protein structure was represented as a graph based on residue interactions, where nodes correspond to residues and edges denote non-covalent interactions between them. The Solvent Accessible Surface Area (SASA) was calculated [28], and residues exhibiting an exposure of 23.5% or greater were classified as surface residues, following the threshold suggested in [35, 48]. The final graph comprises only surface residues together with their first two neighbouring shells. These graphs thus capture spatial and topological context essential for interaction modelling. Figure 5 illustrates a graph representation of a protein structure as described.

**Fig. 5.**
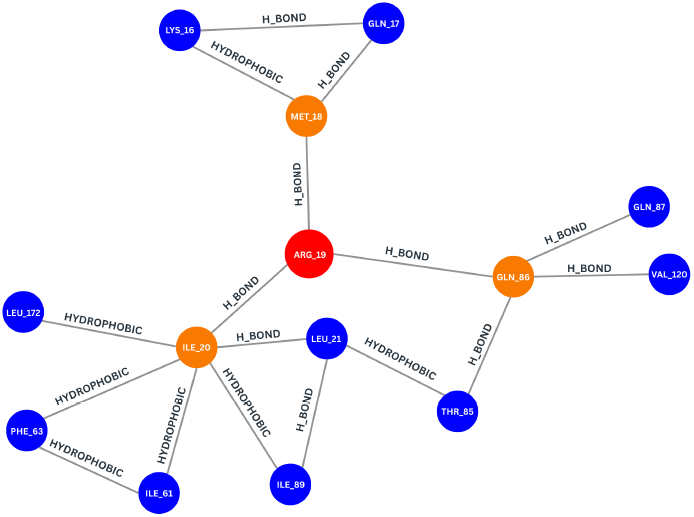
Subgraph representing a surface residue and its first two neighbouring shells for the AlphaFold model AF-P84085-F1 (chain A). The subgraph is centred on the surface residue ARG:19. The red node denotes the surface residue of interest; gold-coloured nodes represent adjacent residues (1^st^ shell), and blue nodes correspond to adjacent–adjacent residues (2^nd^ shell). Edges indicate interaction types, such as hydrogen bonds and hydrophobic interactions. The diagram highlights the multi-layer connectivity and interaction topology surrounding the surface residue.

Protein graphs are encoded as feature vectors, including features of surface residues and their first and second shell neighbors. The features comprise residues (the 20 standard ones), SASA value, hydrophobicity, charge, number of SASA residues, 3D coordinates of the alpha carbon, atomic types, and interaction types. A summarized breakdown of these feature vectors is provided in Figure S4 of the Supplementary Material. A matrix composed of the set of feature vectors was used as input to the PPI type classifier, which employs the Random Forest machine learning algorithm with 10-fold cross-validation. Overall evaluation metrics are 57.74% accuracy, 57.54% precision, 57.74% recall and 57.45% F1 score. Table 2 presents the model’s performance detailed metrics across all the interaction types.

**Table 2.**
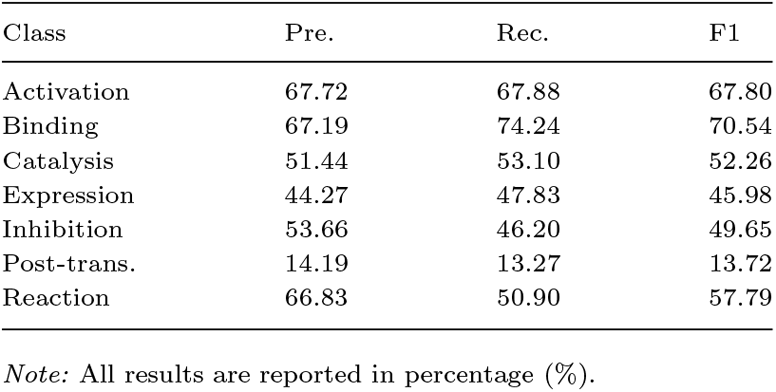
Class-wise Performance Metrics.

We compared the results with existing methods SCNN, Mine, PIPR, TAGPPI. Our method outperforms previous ones in predicting PPI types, indicating that physicochemical and topological features used as descriptors, derived from surface residues, can effectively represent proteins. Table 3 summarizes the model’s comparison with competitors.

**Table 3.**
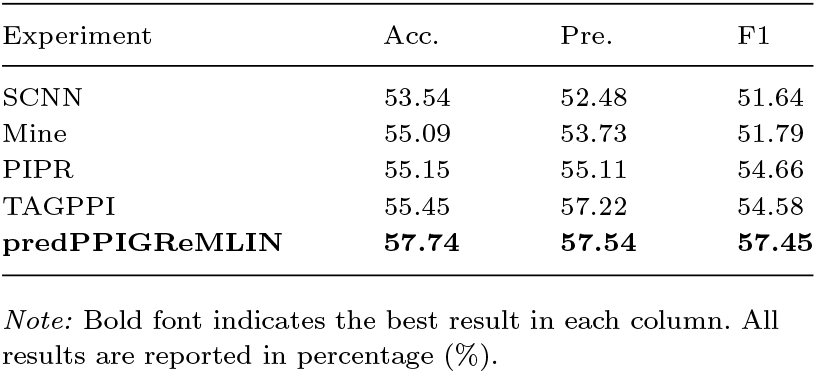
Comparison of Model Performance.

### Case study: predicting ligands for SARS-CoV-2 RBD

To validate our predictions, we performed docking simulations for the ligands in the SARS-CoV-2 spike–ACE2 pockets using HADDOCK3. We compared the binding energies of the predicted ligands with those of real ligands through a redocking protocol. The hypothesis was that our proposed method could predict a ligand that interacts with the target protein, maintaining the binding energies features of the experimental complexes. Specifically, we aimed to determine whether our method could predict ligands able to bind the target, forming complexes that retain the key characteristics of the SARS-CoV-2 RBD interaction. Additionally, to assess the statistical significance of our results, we compared them with the null ligands. The null ligands were selected at random from PDB protein-protein complexes. We focused on the mean binding free energy, its statistical dispersion and probability density to gain insights into the predictive power of our method. Table 4 summarizes the statistical analysis of the binding free energy.

**Table 4.**
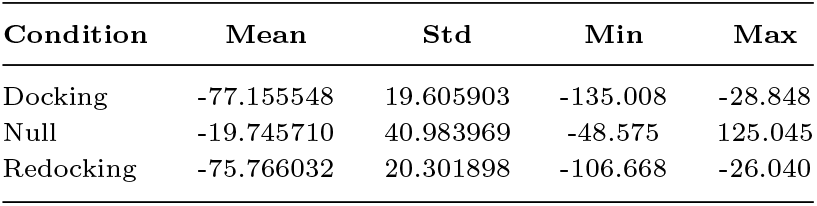
Statistical Analysis of HADDOCK Scores.

The evaluation revealed that the mean binding energies score for the *docking* group (−77.16) closely matched that of the *redocking* reference (−75.77). This suggests that the complexes with ligands predicted by predPPIGReMLIN successfully replicated the energetic characteristics of the spike–ACE2 complex. In contrast, the complexes involving null ligands showed markedly higher (less negative) mean scores (−19.75), reflecting reduced stability and poor interface complementarity.

To quantitatively assess the similarity of the binding energies scoring profiles between the docking scenarios, we compared the distributions of binding free energies. A two-sample t-test was used to evaluate the differences in mean binding energies scores between the Docking, Redocking, and Null datasets.

Figure 6 illustrates that the binding energy distribution for the Docking closely matches that of the Redocking energies. This suggests that the predicted ligands exhibit similar binding characteristics to those observed in experimentally validated complexes. In contrast, the distribution of the Null ligands is markedly different from both the Docking and Redocking distributions, indicating that the predicted ligands are energetically more favorable compared to random, non-specific ligands.

**Fig. 6.**
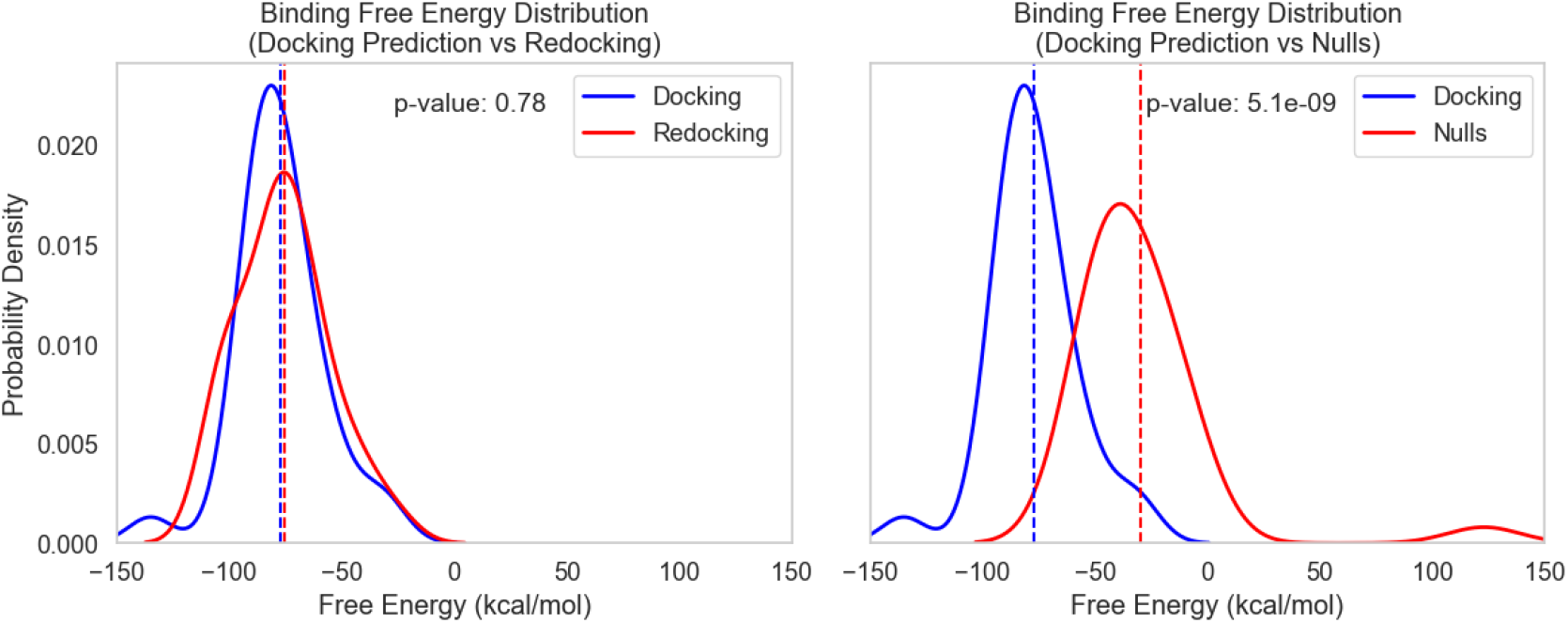
Comparison of HADDOCK binding free energy distributions between Docking, Redocking, and Null complexes. Kernel density estimates (KDEs) depict the probability distribution of HADDOCK-derived binding free energies. The **(Left)** panel shows the Docking (blue) and Redocking (red) energy profiles, while the **(Right)** panel compares Docking and Null datasets. Dashed vertical lines indicate mean values for each distribution. Statistical significance was evaluated using Welch’s two-sample t-test. No significant difference was observed between Docking and Redocking (*p* = 0.78), whereas Docking and Null distributions exhibited a highly significant difference (*p* = 5.1 × 10^−9^).

T-tests were conducted to compare the mean binding energies between the Docking and Redocking groups, yielding a high p-value of (*p* = 0.78), which indicates no significant difference between these two groups. However, the comparison between Docking and Null groups produced a very low p-value (*p* = 5.1 × 10^−9^), demonstrating a statistically significant difference. This suggests that the interactions established by ligands predicted by predPPI-GReMLIN are energetically favorable and biologically relevant, distinguishing them from interactions formed by random ligands.

Collectively, the comparison of the binding free energy distributions reveals that the predicted ligands exhibit binding profiles similar to those obtained from redocking, while being significantly different from random ligands. This evidence suggests that predPPI-GReMLIN can predict meaningful and energetically favorable ligands, capable to establish protein–protein interactions.

## Conclusion

In this study, we introduced predPPI-GReMLIN, a graph-based computational framework designed to predict protein–protein interactions (PPIs) by integrating atomic-level structural and physicochemical features with conserved subgraph mining. Unlike sequence-based approaches that rely primarily on amino acid order and co-evolutionary signals, our model captures the intrinsic three-dimensional nature of molecular recognition by representing protein–protein interfaces as bipartite interaction graphs. Nodes denote atoms and edges encode non-covalent interactions governed by spatial distance and physicochemical compatibility.

The proposed method extends the original ppiGReMLIN algorithm beyond structural motif identification to ligand and interaction prediction, through a novel graph search and ligand-swap mechanism. This design enables the identification of potential binding partners by searching a large graph corpus derived from experimentally solved protein complexes and selecting pattern-matched ligands based on conserved interfaces.

In the evaluation of binary interaction prediction tasks using the CAMP and Yeast datasets, predPPI-GReMLIN achieved accuracy, precision, recall, and F1-scores exceeding 97%, outperforming all evaluated state-of-the-art sequence-based models, including TAGPPI, PIPR, and DeepPPI. The high MCC further attests to the model’s discriminative capacity and stability.

In the classification of protein interaction types, the use of solvent accessible surface area (SASA) features to model surface residue exposure and local environment yielded a 57.74% classification accuracy, outperforming competing methods such as SCNN, Mine, and TAGPPI. This improvement highlights the contribution of physicochemical descriptors and spatial topology in capturing the subtle determinants of diverse interaction modalities, ranging from catalytic to inhibitory or signaling relationships.

A case study on the SARS-CoV-2 spike–ACE2 complex further illustrated the predictive ability of the model. The docking with predicted ligands generated complexes whose HADDOCK3 scores closely mirrored native redocking results. The docking and redocking groups exhibited overlapping low-energy distributions, while the docking with null ligands displayed higher and more variable scores. The result confirms the predPPI-GReMLIN’s ability to predict interacting partners the form complexes that reproduce native-like binding energetics and structural complementarity.

Overall, the findings highlight that predPPI-GReMLIN is a scalable, interpretable, and biologically grounded method for PPI analysis. Its reliance on conserved subgraph motifs, combined with prediction of ligands, makes it suitable not only for understanding known interactions but also for discovering novel protein partners and guiding rational drug or peptide design. Future extensions may include the integration of graph neural networks (GNNs) to learn hierarchical structural embeddings, the incorporation of dynamic conformational states from molecular simulations, and the expansion of the graph corpus to use protein models.

## Appendix: Supplementary Materials

Additional data supporting this study are provided in the supplementary document. The datasets and source code used in this study are available at: https://github.com/morufwork/predPPIGReMLIN.git

## Competing interests

No competing interest is declared.

## Acknowledgments

This study was financed in part by the Coordenação de Aperfeiçoamento de Pessoal de Nível Superior - Brasil (CAPES); Conselho Nacional de Desenvolvimento Científico e Tecnológico (CNPq) under grant number 444426/2024-8; Fundação de Amparo à Pesquisa do Estado de Minas Gerais (FAPEMIG). The funding agencies had no role in study design, data collection, analysis and interpretation, decision to publish, or preparation of the manuscript.

## Notes

### Competing Interest Statement

The authors have declared no competing interest.

