## Supplementary material for "predPPI-GReMLIN: prediction of protein-protein interactions through mining of conserved bipartite graphs": Table S1 and S2, Figure S1, s2, s3, and S4

### FIGURES

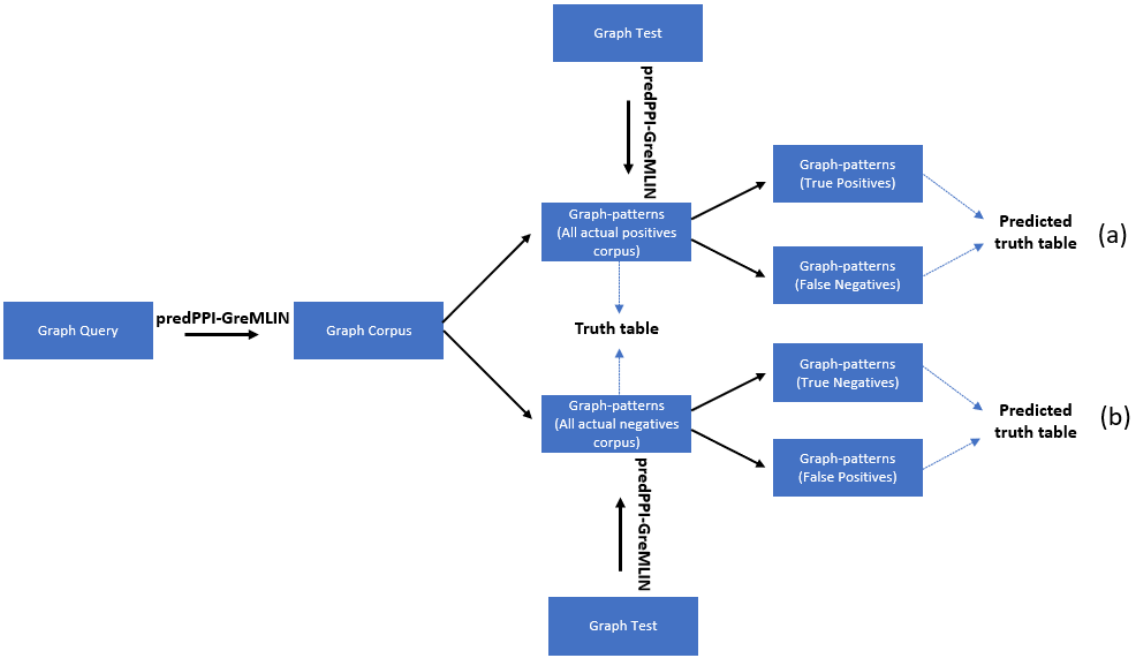


**Figure S1.** Construction of the prediction truth table using graph-pattern matching in predPPI-GReMLIN. Through predPPI-GReMLIN, the corpus is divided into two subsets: Graph-patterns of all actual positives (binary 1), representing known interacting pairs, and Graph-patterns of all actual negatives (binary 0), representing non-interacting pairs. The truth table was constructed using the concept of a confusion matrix, comparing test-derived patterns against these two corpora to assess the model's prediction accuracy. The concept is as follows: **(a)** All actual positives = True Positive (TP) + False Negative (FN); **(b)** All actual negatives = True Negative (TN) + False Positive (FP). This enabled a binary classification of predicted versus actual graph-pattern matches, forming the basis for performance evaluation. Positive Graph Pattern Matching: Graph-based pattern matching was executed between graph test T(G) and all actual positive graph patterns. The results were compiled into a prediction truth table with TP and FN classifications. Negative Graph Pattern Matching: A similar process was performed between graph test T(G) and all actual negative graph patterns, generating TN and FP classifications.

| (a) | Graph level features | | | | | | | | | | Centrality features | | | | Node metadata aggregation | | | | | | Edge attribute aggregation | | | | | |
|---|---|---|---|---|---|---|---|---|---|---|---|---|---|---|---|---|---|---|---|---|---|---|---|---|---|---|
| (b) | NN | NE | DS | CS | ACC | TS | AT | MaD | MiD | AD | AB | AC | DM | RD | AL | UR | AA | --- | NT | --- | AED | MiED | MaED | ET | --- | Class |
| (c) | 2 | 1 | 1 | 1 | 0 | 0 | 0 | 1 | 1 | 1 | 0 | 1 | 1 | 1 | 0.5 | 2 | 0 | --- | 1 | --- | 2.81 | 2.81 | 2.81 | 0 | --- | 1 |
| | 14 | 22 | 0.242 | 1 | 0 | 0 | -0.68 | 8 | 1 | 3.143 | 0.098 | 0.474 | 4 | 2 | 0.286 | 4 | 0 | --- | 0 | --- | 4.31 | 2.92 | 5.86 | 0 | --- | 0 |
| | 6 | 5 | 0.333 | 1 | 0 | 0 | -0.25 | 2 | 1 | 1.667 | 0.333 | 0.448 | 5 | 3 | 0.5 | 2 | 0 | --- | 0 | --- | 3.72 | 3.54 | 3.78 | 0 | --- | 1 |
| | 9 | 14 | 0.389 | 1 | 0 | 0 | -0.75 | 6 | 2 | 3.111 | 0.103 | 0.590 | 3 | 2 | 0.333 | 3 | 0 | --- | 0 | --- | 4.14 | 2.52 | 5.86 | 0 | --- | 0 |

**Figure S2.** prePPi-GReMLIN feature vectors for binary classification of graph patterns. **(a)** Starting top-down, the first row consists of four high-level columns, each with a column group of graph-level features, centrality features, node metadata aggregation, and edge attribute aggregation. **(b)** The second row is segmented in columns: in the graph level features, we have number of nodes (NN), number of edges (NE), density (DS), connectivity status (CS), average clustering coefficient (ACC), transitivity (TS), assortativity (AT), max degree (MaD), min degree (MiD), and average degree (AD); in the centrality features, we have average betweenness (AB), average closeness (AC), diameter (DM), and radius (RD); in the node metadata aggregation, we have average ligand presence (AL), number of unique residues (UR), 20 standard amino acid identifiers (AA), and 19 node type count (NT); and in the edge attribute aggregation, we have average edge distances (AED), minimum edge distances (MiED), and maximum edge distances (MaED), and 11 edge type counts (ET). We have a total of 69 features across all four high-level columns. The last column represents interacting patterns or no interacting patterns (class). **(c)** Hereafter, each row corresponds to a graph, and each column corresponds to a descriptor and a class.

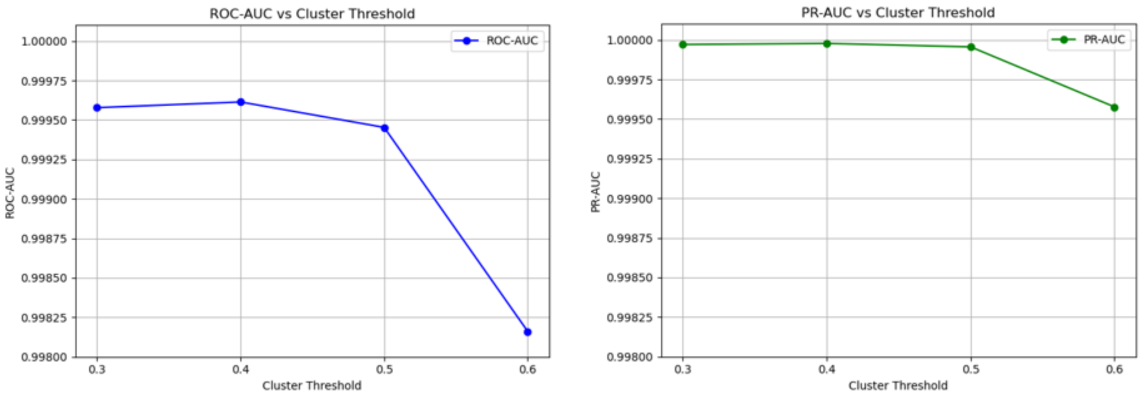


**Figure S3.** How clustering threshold affects model performance. **(Left):** The ROC-AUC scores for the Random Forest classifier across four clustering thresholds (0.3, 0.4, 0.5, 0.6). A slight decrease in ROC-AUC is observed as the threshold increases, particularly at threshold 0.6. **(Right):** The corresponding PR-AUC scores remain nearly perfect up to threshold 0.5, after which a slight drop is observed at 0.6. These results indicate that the model generalizes well across different granular clusters, but extremely fine-grained clusters may introduce noise or instability due to smaller cluster sizes.

| (a) | P1 | | | | | | | | | | | | | | | | | | | | | | | | P2 | | |
|---|---|---|---|---|---|---|---|---|---|---|---|---|---|---|---|---|---|---|---|---|---|---|---|---|---|---|---|
| (b) | Surface residue | | | | | | | | | | | | | | | | | | | First shell | | | Second Shell | | | | |
| (c) | Residue level properties | | | | | | | Res. | Atoms | | | | | | Interactions | | | | | Res. | Atoms | | Res. | | | | |
| (d) | **AA** | **---** | **SA** | **HYP** | **CH** | **CD** | **---** | **RC** | **ACP** | **DON** | **POS** | **NEG** | **HPB** | **ARM** | **HY** | **SB** | **AS** | **HB** | **RP** | **RC** | **ACP** | **---** | **RC** | **---** | **---** | **---** | **Class** |
| (e) | 0.09 | --- | 90.7 | -0.55 | -0.08 | 6.71 | --- | 515 | 868 | 735 | 196 | 228 | 927 | 157 | 51 | 45 | 0 | 1008 | 24 | 515 | 928 | --- | 2236 | --- | --- | --- | 6 |
| | 0.04 | --- | 57.6 | -0.12 | -0.01 | -6.08 | --- | 349 | 582 | 597 | 202 | 110 | 792 | 251 | 50 | 27 | 1 | 882 | 14 | 349 | 496 | --- | 2016 | --- | --- | --- | 1 |
| | 0.07 | --- | 43.5 | -0.18 | 0.001 | 1.51 | --- | 254 | 442 | 449 | 189 | 120 | 569 | 167 | 54 | 64 | 2 | 628 | 25 | 254 | 404 | --- | 1573 | --- | --- | --- | 1 |
| | 0.07 | --- | 77.0 | -0.35 | -0.06 | 12.3 | --- | 474 | 782 | 705 | 194 | 172 | 1056 | 250 | 60 | 32 | 0 | 970 | 30 | 474 | 642 | --- | 2092 | --- | --- | --- | 6 |
| | 0.04 | --- | 67.0 | 0.13 | -0.02 | 2.16 | --- | 93 | 147 | 139 | 26 | 22 | 228 | 107 | 17 | 4 | 3 | 220 | 2 | 93 | 146 | --- | 516 | --- | --- | --- | 6 |

**Figure S4.** predPPI-GReMLIN feature vectors. **(a)** Starting top-down, the first row consists of a pair of proteins whose feature vectors are considered. **(b)** In the second row, the first row consists of three high-level columns: the surface residue, the first and the second shell of neighbors, respectively. **(c)** In the third row, each column groups descriptors at the residue, interaction, and atom levels for each high-level column mentioned in (b). **(d)** The fourth row is segmented in columns: in residue level properties, we have mean of each 20 standard amino acid identifiers (AA), solvent accessibility values (SA), hydrophobicity (HYP), charge (CH), and the 3D coordinates of the Cα atom (CD); In the residue, we have residue count (RC); In the atom level, we have acceptor (ACP), donor (DON), positive (POS), negative (NEG), hydrophobic (HPB), and aromatic (ARM); and In the interaction level, we have hydrophobic (HY), salt bridge (SB), aromatic stacking (AS), hydrogen bond (HB), and repulsive (RP). We have only the residue count, atomic types, and interaction type descriptors for the first and second shells, totaling 62 descriptors per protein structure and 124 for a pair (P1, P2) of protein structures. The last column represents the types of interaction between a pair of protein structures. € Hereafter, each row represents a pair of protein structures, and each column represents a residue descriptor and corresponding interaction types.

### TABLES

In Table S1, to assess the performance and generalization capability of the proposed predPPI-GReMLIN model, a five-fold cross-validation strategy was employed. In this evaluation, the dataset was randomly divided into five equal-sized subsets, with four used for training and one for testing in each fold. The process was repeated five times to ensure robustness, with each subset serving as the test set once. The averaged performance metrics—accuracy, precision, recall, F1-score, and AUC—provide a comprehensive evaluation of the model's predictive ability across different partitions of the dataset. Table S1 presents the overall evaluation of predPPI-GReMLIN across five-fold cross-validation.

**Table S1.** Overall evaluation of the predPPI-GReMLIN across five folds of cross-validation

| **F1 Fold** | | | | | | | | | | |
|---|---|---|---|---|---|---|---|---|---|---|
| | TP | FN | TN | FP | Accuracy | Precision | Sensitivity | Specificity | F1 Score | MCC |
| F2 on F1 | 108775 | 450 | 5553 | 447 | 0.992215 | 0.995907 | 0.99588 | 0.9255 | 0.995894 | 0.921162 |
| F3 on F1 | 108889 | 336 | 5545 | 455 | 0.993135 | 0.995839 | 0.996924 | 0.924167 | 0.996381 | 0.929856 |
| F4 on F1 | 108906 | 319 | 5505 | 495 | 0.992936 | 0.995475 | 0.997079 | 0.9175 | 0.996277 | 0.927549 |
| F5 on F1 | 108867 | 358 | 5510 | 490 | 0.99264 | 0.995519 | 0.996722 | 0.918333 | 0.99612 | 0.924732 |
| | | | | **Mean** | **0.992732** | **0.995685** | **0.996651** | **0.921375** | **0.996168** | **0.925825** |
| **F2 Fold** | | | | | | | | | | |
| F1 on F2 | 108775 | 447 | 5553 | 450 | 0.992215 | 0.99588 | 0.995907 | 0.925037 | 0.995894 | 0.921162 |
| F3 on F2 | 108896 | 326 | 5555 | 448 | 0.993283 | 0.995903 | 0.997015 | 0.925371 | 0.996459 | 0.931384 |
| F4 on F2 | 108904 | 318 | 5506 | 497 | 0.992927 | 0.995457 | 0.997088 | 0.917208 | 0.996272 | 0.927481 |
| F5 on F2 | 108889 | 333 | 5535 | 468 | 0.993048 | 0.99572 | 0.996951 | 0.922039 | 0.996335 | 0.928927 |
| | | | | **Mean** | **0.992868** | **0.99574** | **0.996741** | **0.922414** | **0.99624** | **0.927239** |
| **F3 Fold** | | | | | | | | | | |
| F1 on F3 | 108889 | 455 | 5545 | 336 | 0.993135 | 0.996924 | 0.995839 | 0.942867 | 0.996381 | 0.929856 |
| F2 on F3 | 108896 | 448 | 5555 | 326 | 0.993283 | 0.997015 | 0.995903 | 0.944567 | 0.996459 | 0.931384 |
| F4 on F3 | 108977 | 367 | 5457 | 424 | 0.993135 | 0.996124 | 0.996644 | 0.927903 | 0.996384 | 0.928818 |
| F5 on F3 | 109017 | 327 | 5541 | 340 | 0.994211 | 0.996891 | 0.997009 | 0.942187 | 0.99695 | 0.94018 |
| | | | | **Mean** | **0.993441** | **0.996739** | **0.996349** | **0.939381** | **0.996543** | **0.93256** |
| **F4 Fold** | | | | | | | | | | |
| F1 on F4 | 108906 | 495 | 5505 | 319 | 0.992936 | 0.997079 | 0.995475 | 0.945227 | 0.996277 | 0.927549 |
| F2 on F4 | 108904 | 497 | 5506 | 318 | 0.992927 | 0.997088 | 0.995457 | 0.945398 | 0.996272 | 0.927481 |
| F3 on F4 | 108977 | 424 | 5457 | 367 | 0.993135 | 0.996644 | 0.996124 | 0.936985 | 0.996384 | 0.928818 |
| F5 on F4 | 109013 | 388 | 5480 | 344 | 0.993647 | 0.996854 | 0.996453 | 0.940934 | 0.996654 | 0.934054 |
| | | | | **Mean** | **0.993161** | **0.996916** | **0.995878** | **0.942136** | **0.996397** | **0.929476** |
| **F5 Fold** | | | | | | | | | | |
| F1 on F5 | 108867 | 490 | 5510 | 358 | 0.99264 | 0.996722 | 0.995519 | 0.938991 | 0.99612 | 0.924732 |
| F2 on F5 | 108889 | 468 | 5535 | 333 | 0.993048 | 0.996951 | 0.99572 | 0.943252 | 0.996335 | 0.928927 |
| F3 on F5 | 109017 | 340 | 5541 | 327 | 0.994211 | 0.997009 | 0.996891 | 0.944274 | 0.99695 | 0.94018 |
| F4 on F5 | 109013 | 344 | 5480 | 388 | 0.993647 | 0.996453 | 0.996854 | 0.933879 | 0.996654 | 0.934054 |
| | | | | **Mean** | **0.993387** | **0.996784** | **0.996246** | **0.940099** | **0.996515** | **0.931973** |
| | | | | **Result** | **0.993118** | **0.996373** | **0.996373** | **0.933081** | **0.996373** | **0.929414** |
| | | | | **Final Result** | **99.31** | **99.64** | **99.64** | **93.31** | **99.64** | **92.94** |

In Table S2, the protein–protein interaction dataset contains pairs of interacting human proteins represented by their Ensembl Protein (ENSP) identifiers. Each record (P1, P2) corresponds to a distinct protein interaction, annotated with a class label ranging from 0 to 6, representing different interaction types. Protein identifiers and sequences were first linked to UniProt (UP) identifiers. The UniProt identifiers were used to retrieve AlphaFold identifiers, which were subsequently used to obtain the corresponding AlphaFold-predicted protein structures from the AlphaFold database for subsequent modeling. This curated dataset serves as the foundation for analyzing structural features and predicting interaction outcomes using graph-based methods**.** Table S2 summarizes the mapping between UniProt and AlphaFold identifiers.

**Table S2.** Mapping Between UniProt and AlphaFold IDs

| **P1 (Protein 1)** | **P2 (Protein 2)** | **Class Label** | **UniProt ID (P1)** | **UniProt ID (P2)** | **AlphaFold ID (P1)** | **AlphaFold ID (P2)** |
|---|---|---|---|---|---|---|
| 9606.ENSP00000019103 | 9606.ENSP00000275216 | 2 | P47872 | Q96RJ0 | AF-P47872-F1-model_v4.pdb | AF-Q96RJ0-F1-model_v4.pdb |
| 9606.ENSP00000215832 | 9606.ENSP00000267396 | 2 | P28482 | Q8IYK8 | AF-P28482-F1-model_v4.pdb | AF-Q8IYK8-F1-model_v4.pdb |
| 9606.ENSP00000216190 | 9606.ENSP00000248342 | 5 | O15371 | Q9UBQ5 | AF-O15371-F1-model_v4.pdb | AF-Q9UBQ5-F1-model_v4.pdb |
| 9606.ENSP00000232461 | 9606.ENSP00000263125 | 0 | P11488 | Q04759 | AF-P11488-F1-model_v4.pdb | AF-Q04759-F1-model_v4.pdb |
